# Local Monomer Levels and Established Filaments Potentiate Non-Muscle Myosin 2 Assembly

**DOI:** 10.1101/2023.04.26.538303

**Authors:** Melissa A. Quintanilla, Hiral Patel, Huini Wu, Kem A. Sochacki, Matthew Akamatsu, Jeremy D. Rotty, Farida Korobova, James E. Bear, Justin W. Taraska, Patrick W. Oakes, Jordan R. Beach

**Affiliations:** Department of Cell and Molecular Physiology, Stritch School of Medicine, Loyola University Chicago, Maywood, IL; Laboratory of Molecular Biophysics, National Heart, Lung, and Blood Institute, NIH, Bethesda, MD, USA; Department of Biology, University of Washington, Seattle, WA; Department of Biochemistry, Uniformed Services University of the Health Sciences, Bethesda, MD; Feinberg School of Medicine, Northwestern University, Chicago, IL; Department of Cell Biology and Physiology, University of North Carolina-Chapel Hill, Chapel Hill, NC

**Keywords:** Non-muscle myosin 2, assembly, amplification, molecular counting, optogenetics, CLEM, actin

## Abstract

The ability to dynamically assemble contractile networks is required throughout cell physiology, yet the biophysical mechanisms regulating non-muscle myosin 2 filament assembly in living cells are lacking. Here we use a suite of dynamic, quantitative imaging approaches to identify deterministic factors that drive myosin filament appearance and amplification. We find that actin dynamics regulate myosin assembly, but that the actin architecture plays a minimal direct role. Instead, remodeling of actin networks modulates the local myosin monomer levels and facilitates assembly through myosin:myosin driven interactions. Using optogenetically controlled myosin, we demonstrate that locally concentrating myosin is sufficient to both form filaments and jump-start filament amplification and partitioning. By counting myosin monomers within filaments, we demonstrate a myosin-facilitated assembly process that establishes sub-resolution filament stacks prior to partitioning into clusters that feed higher-order networks.

Together these findings establish the biophysical mechanisms regulating the assembly of non-muscle contractile structures that are ubiquitous throughout cell biology.

## Introduction

Non-muscle myosin 2 (NM2) is a cytoskeletal motor protein that builds bipolar filaments to engage actin filaments and generate contractile forces. The magnitude and orientation of these forces are highly tunable to regulate processes at the cell, tissue, and organism level (1). This adaptability across spatial and temporal scales requires active remodeling of actomyosin networks. Understanding the spatiotemporal mechanisms of how cells build these force-producing units is therefore critical.

NM2 filaments are dynamically assembled from NM2 monomers, which consist of two myosin heavy chains (MHC), two essential light chains (ELC), and two regulatory light chains (RLC). Each MHC consists of an N-terminal motor domain, light chain-binding neck region, and a C-terminal alpha helix which dimerizes into a coiled-coil tail. The standard monomer-to-filament model of NM2 filament assembly begins with phosphorylation of RLC on Thr18/Ser19 (2), which drives the NM2 monomer from the folded, inactive 10S state to the unfolded, assembly-competent 6S state (3, 4). Once unfolded, the coiled-coil tails readily associate in parallel and anti-parallel orientations to form a bipolar filament (5). Kinases from a variety of signaling networks phosphorylate the RLC to enhance NM2 filament assembly, with the dominant kinases being RhoA-activated Rho-associated coiled-coil kinase (ROCK1/2) and Ca^++^/calmodulin-activated NM2 light chain kinase (MLCK) (6). In addition to phospho-modulation, *in vitro* studies demonstrated that NM2 filament assembly was enhanced in the presence of actin filaments (7), suggesting combinatorial contributions from both kinase signaling and actin networks.

To explore molecular details of NM2 filament assembly in living cells, it is important to capture data at the length and time scales of the interactions in question. Recent advances in light microscopy have provided the spatial resolution required to observe discrete NM2 filaments (*∼* 300 nm in length) with the temporal resolution required to observe network assembly (8, 9). These studies have added dynamic mechanistic insight to earlier static electron microscopy (EM) experiments (10, 11), and demonstrated that the simple monomer-to-filament model is incomplete in cellular contexts. More specifically, we and others observed that once an initial NM2 filament is established by unknown mechanisms in the lamella of a migrating cell, it grows in intensity, and then “partitions” into a cluster of filaments or “expands” into a stack of filaments (8, 9). These clusters/stacks then merge with the higher-order actomyosin networks within the cell (stress fibers, transverse arcs, etc.). Similar progressions have been observed in contractile ring assembly, suggesting a common and universal mechanism for initiating and amplifying NM2 networks (12).

Despite these technology-enabled advances, we currently lack an experimentally-supported working model for how a nascent NM2 filament is precisely established in space and time within a cell. We also do not understand how nascent NM2 filaments contribute to the higher-order network assembly required for physiological levels of contraction. Here we show that leading edge retractions are better predictors of NM2 filament assembly than canonically proposed calcium and RhoA signaling events. Similarly, we find that actin dynamics regulate NM2 filament assembly, decreasing assembly when actin dynamics are stalled, and amplifying assembly following the break down of actomyosin structures elsewhere in the cell. Despite the clear role for actin dynamics in NM2 assembly, we did not observe an actin ultrastructure that is prognostic of NM2 filament formation. Instead, we find that by locally increasing myosin concentration, we can assemble NM2 filaments and initiate filament amplification and further partitioning. Finally, using molecular standard candles, we count the number of myosin monomers in filaments and show that monomers are more likely to add to existing myosin clusters instead of forming nascent filaments. We also find that partitioning myosin typically already contain multiple filaments, suggesting that amplification precedes partitioning. Together these findings clarify the dynamics of NM2 filament assembly within cells.

## Results

### Leading edge retractions precede nascent NM2 filament appearance

To better understand the precise events that precede nascent NM2 filament assembly in cells, we initially tested spatiotemporal correlation of filament appearance with known upstream biochemical modulators.

We generated fibroblasts (13) with the N-terminus of endogenous NM2A tagged with mScarlet (mScarlet-NM2A), and exogenously expressed established fluorescent biosensors to localize calcium (GCaMP7s) or active RhoA (Anillin AHPH) (14, 15). We then imaged the lamella of migrating cells, where discrete NM2 assembly events can readily be observed with high resolution light microscopy (Fig.1A, Movie 1) (9).

**Fig. 1.**
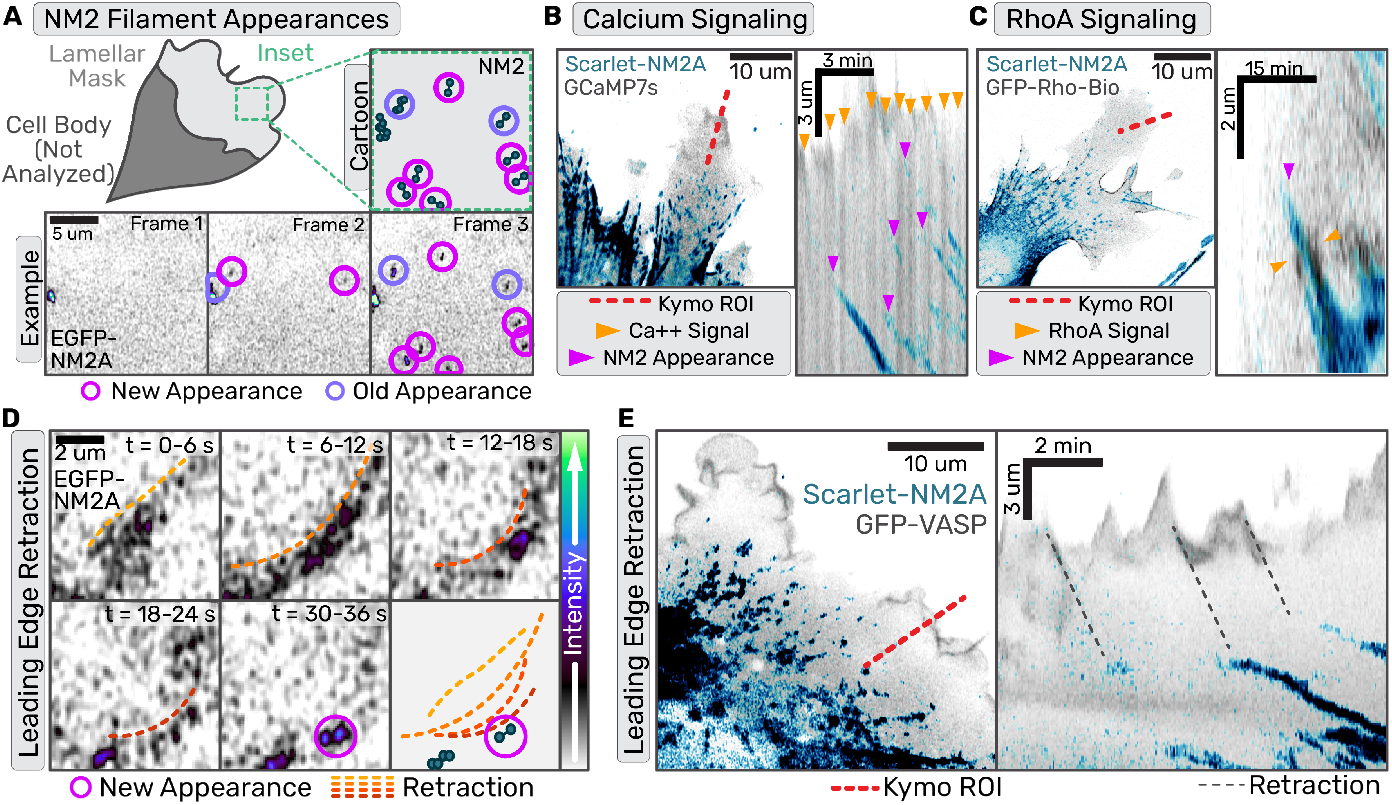
Nascent NM2 filament appearances correlate with leading edge retraction events but not upstream signaling. (a,d,e) Confocal z-stacks of primary EGFP-NM2A MEF cells. (b,c) Confocal z-stacks of immortalized Scarlet-NM2A MEF knockin cells. (a) Example of NM2 filament appearance detection workflow with both cartoon and example frames. (b) Scarlet-NM2A (in blue) and GCaMP7s (in grey) imaged with Zeiss Airyscan 880. Left panel displays example frame with kymograph ROI indicated by red dotted line. In the corresponding kymograph in the right panel, orange arrows mark calcium sparks and magenta arrows mark NM2 filament appearance. Scale bar= 10 um.(c) Scarlet-NM2A (in blue) and RhoA biosensor (in grey) imaged with Zeiss Airyscan 880. Left panel displays example frame with kymograph ROI indicated by red dotted line. In the corresponding kymograph in the right panel, orange arrows mark active RhoA signal and magenta arrows mark NM2 filament appearance. Scale bar = 10 um. (d) Sum intensity projection of z-stack and 3 frame time averaging for frames collected every second. Orange-to-red gradient dotted lines indicate wave-like retraction and magenta circles indicate subsequent NM2 filament appearance. Scale bar = 2 um. (e) Scarlet-NM2A cells transiently expressing mEGFP-VASP imaged with Zeiss 880 confocal every second. Image is a sum intensity projection of Z-stack with NM2 in blue and vasp in purple. Scale bar = 10 um. Red line indicates the ROI used for subsequent kymograph in right panel. Grey dotted lines connect leading edge retractions with NM2 filament appearance.

Although we detected both calcium and RhoA activity in the lamella, neither signaling cascade preceded NM2 filament appearance with any apparent precision. When imaging the calcium biosensor, flashes often filled the entire lamella. Occasionally, an NM2 filament appearance followed a calcium flash, but many calcium flashes did not result in filament appearances (Fig.1B). In contrast, active-RhoA did co-localize with NM2 filaments, but after filament appearance instead of preceding it (Fig.1C). Interestingly, active-RhoA often flanked the growing NM2 clusters, reminiscent of NM2-dependent RhoA activation observed in other systems (16). Therefore, while RLC kinases are undoubtedly contributing to NM2 filament assembly, our imaging failed to observe spatiotemporal precision in their contribution to initiating assembly events.

To identify additional factors that might dictate nascent assembly, we assessed lamellar NM2 behavior in polarized fibroblasts. Similar to previous reports (17, 18) we often find NM2 filament appearance is preceded by a leading edge retraction (Fig. 1D, Movie 2). To more carefully observe this correlation, we imaged mScarlet-NM2A fibroblasts expressing EGFP-VASP to visualize the leading edge. Kymographs drawn through the leading edge illustrated multiple retractions of the cell edge that led to the subsequent NM2 filament appearance in the lamella (Fig 1E).

### Actin dynamics facilitate nascent NM2 filament appearance

To directly test the role of leading edge retractions and actin dynamics in NM2 filament assembly, we adopted a drug cocktail consisting of jasplakinolide and latrunculin (JL) that arrests actin dynamics by inhibiting both polymerization and depolymerization (19). First, we confirmed that JL administration stalls leading edge dynamics in the fibroblasts within seconds (Fig 2A-B; Movie 3). We then quantified the rate of NM2 filament appearance in the lamella before and after pharmacological perturbations (Fig 2B-C, Movie 4-5). We found the relative appearance rate did not change in cells treated with DMSO, but significantly decreased upon addition of the actin-stalling JL cocktail (Fig 2C). This demonstrates that while NM2 filament assembly can occur in their absence, leading edge retractions and dynamic actin aid in the process.

**Fig. 2.**
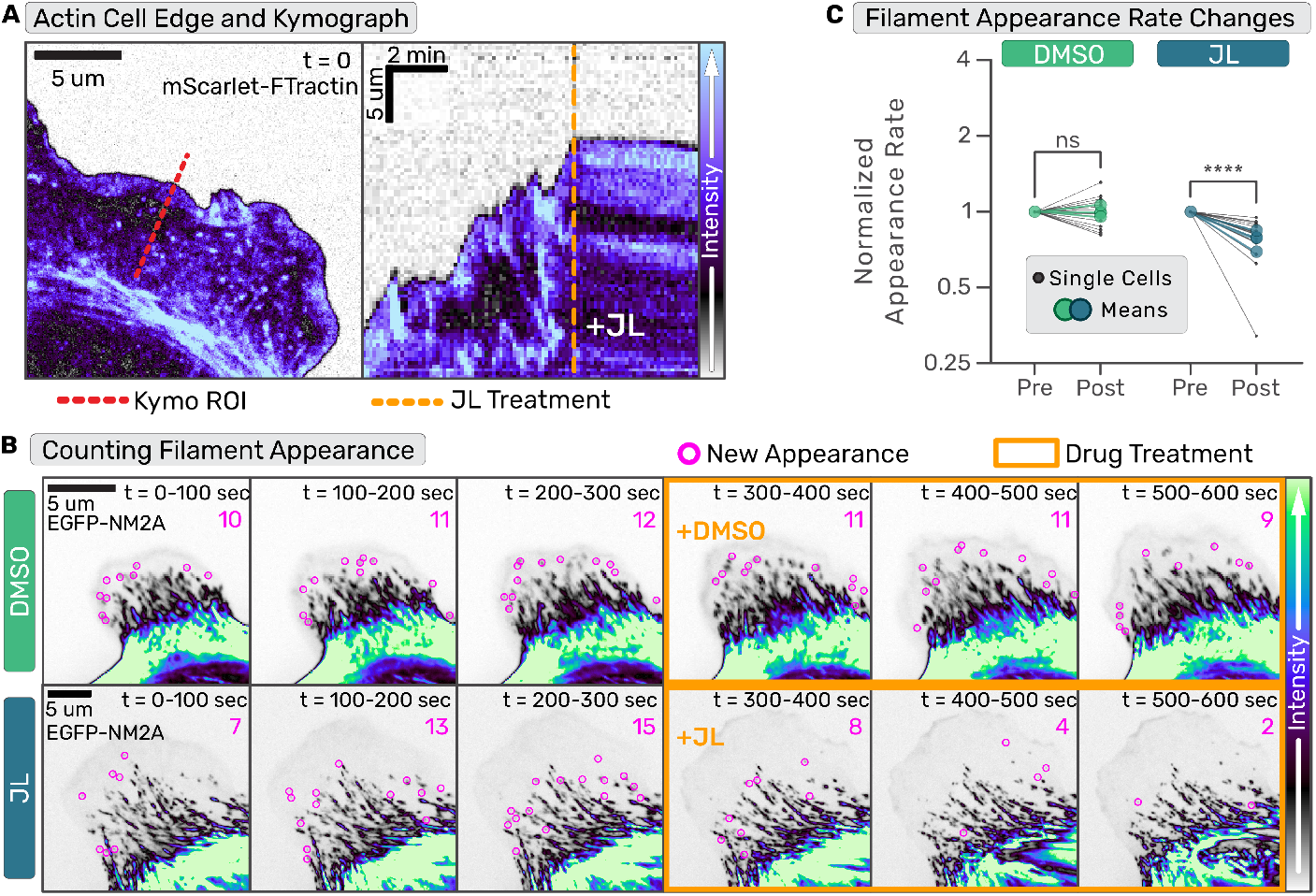
Arresting actin dynamics inhibits nascent NM2 filament appearance. (a-b) Confocal time series of primary EGFP-NM2A MEF cells expressing FTractin-mScarlet were acquired. For (a) only the FTractin-mScarlet channel is shown and for (b) only EGFP-NM2A channel is shown. (a) Example frame of lamellar actin (left panel) and kymograph (right panel) of lamellar dynamics pre- and post-JL treatment through the red dotted line in left panel. Orange dotted line indicates time of JL addition. (b) EGFP-NM2A images were temporally-summed over 20 frames (100 seconds). Nascent NM2 filament appearances during that period are indicated with magenta circles and tallied in upper right corner. Three binned time periods pre- and -post (orange outline) treatment are shown for DMSO (top row) and JL (bottom row). Scale bars = 5 um. (c) Nascent NM2 filament appearance events pre- and post-treatment, normalized to pre-treatment appearance events per minute, were quantified for individual cells (small black dots) and the mean of three independent experiments (large color circles). Temporal and intensity LUTs are indicated to the right. Wilcoxon matched-pairs signed rank test performed comparing each cell (DMSO: 15 cells, JL: 16 cells). p<0.0001

### NM2 filaments assemble in wide array of actin structures

We hypothesized that the underlying actin architecture (alignment, density, etc.) might provide additional cues to facilitate NM2 filament assembly. To better understand the lamellar actin architecture where nascent NM2 filaments are forming, we performed correlative light and platinum replica electron microscopy (PREM; Fig.3) (20, 21). We manually unroofed migrating GFP-NM2A fibroblasts (Fig. 3A) and imaged with both super-resolution fluorescence and platinum replica electron microscopy (Fig 3B-E) (22–24). Within an unroofed lamella, we observe a range of fluorescent NM2A structures, from low intensity doublets with two distinct puncta*∼* 300 nm apart (consistent with a bipolar filament or sub-resolution stack; Fig3F), to larger high intensity clusters with many puncta indicating they contain many NM2A filaments (Fig.3B-C,E). Due to the similar diameter of NM2 bipolar filaments relative to actin filaments, and the overall density of the actin cytoskeleton, we could not distinguish NM2 bipolar filaments in the PREM images, similar to previous reports (11). However, we could observe the local actin architecture where NM2 structures were present and not present. First, the NM2 structures exist in a diverse array of lamellar actin network architectures (Fig. 3D,G). This includes both seemingly disorganized actin and higher density bundled actin (Fig. 3G). Second, while the biggest NM2 clusters typically overlapped with regions of bundled actin, there were no obvious underlying actin features prognostic of low intensity NM2 doublets, with neighboring actin regions appearing indistinguishable from NM2-containing actin regions. Therefore, while filamentous actin supports enhanced NM2 assembly and there are likely structural details with the filamentous actin present here that are beyond the resolution of our PREM imaging, we do not observe specific actin architectures that might be facilitating nascent NM2 assembly events.

**Fig. 3.**
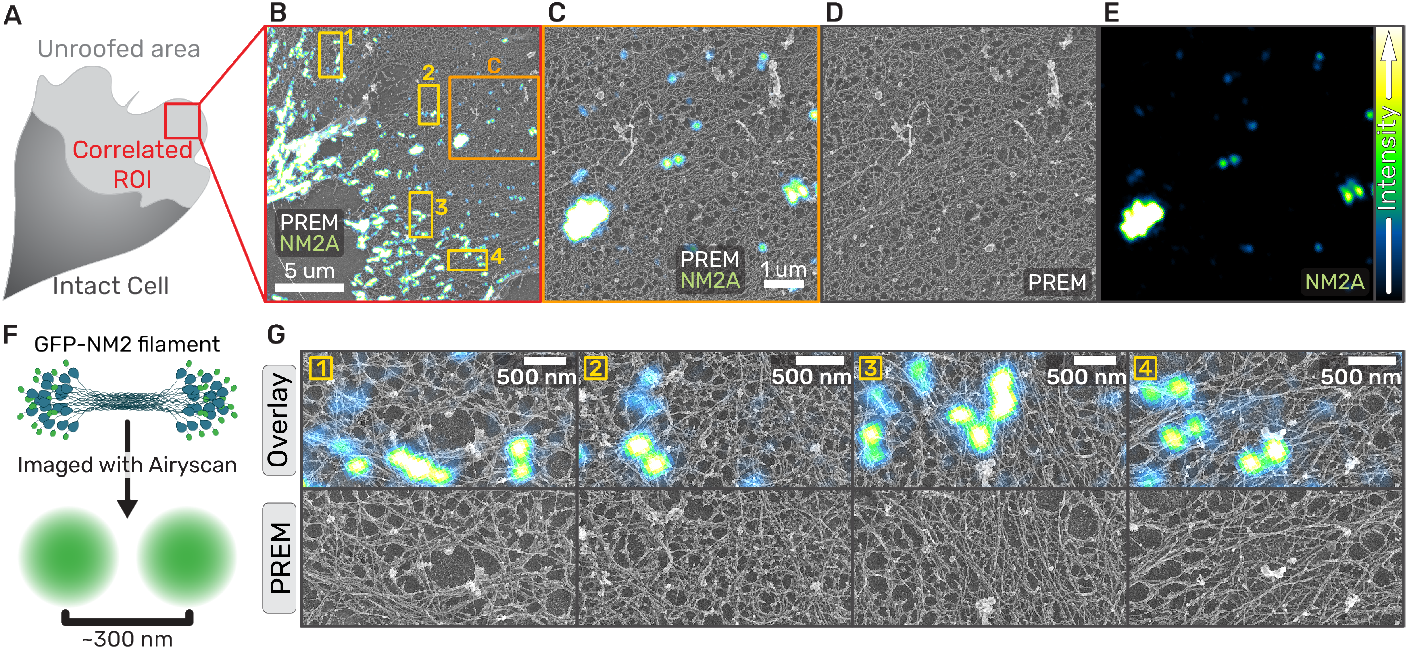
NM2 filaments assemble in wide array of actin architectures. (a) Cartoon of migrating fibroblast with unroofed lamella used for CLEM. (b) Overlay of correlated platinum replica electron micrograph and super-resolution fluorescent micrograph of EGFP-NM2A in an unroofed lamella. Orange box indicates the zoom inset in (c-e) and yellow numbered boxes indicate examples used in (g). (c) Separate images of the zoom inset of the example correlation (left panel) with fluorescent EGFP-2A (right panel) and platinum replica depicting ventral actin network (center panel). Scale bars = 1 um. Intensity LUTs are indicated to the right. (f) Cartoon depicting EGFP-NM2A in filamentous form and the fluorescent signal that is detected from this structure. (g) Examples of EGFP-NM2A clusters in the top ‘overlay’ panel coupled with the underlying actin ultrastructure displayed in the PREM-alone frames in the bottom ‘PREM’ panel. Scale bars = 500 nm.

### Globally elevating NM2 monomer availability initiates filament assembly

Given the lack of an identifying actin ultrastructure to predict NM2 filament formation, we sought to determine whether cytoskeletal dynamics could instead be regulating myosin monomer availability. We used primary MEFs from EGFP-NM2A knockin mice (25) transduced with a lentiviral fluorescent probe for filamentous actin, FTractin-3x-mScarlet (26, 27)), to monitor changes in cell morphology and actin architectures (Movie 6). In long-term, time-lapse imaging of migratory cells we observed a qualitative correlation between NM2 filament appearance and tail retraction events (Fig. 4A, Movie 7). Consistent with previous results (9), we also saw that treatment of cells with ROCK inhibitor (Y27632) resulted not only in disassembly of actomyosin structures, but robust assembly of nascent NM2 filaments in the lamella (Fig. 4B, Movie 8). We therefore hypothesized that global monomer availability in the cytoplasm, whether through changes in morphology or pharmacological perturbation, regulates NM2 filament assembly. To test this hypothesis in the absence of actin dynamics, we treated cells simultaneously with JL and the ROCK inhibitor (JLY; Fig.4C, Movie 9)(19). This JLY treatment not only rescued NM2 filament appearance, but increased it relative to the control (Fig. 4D). This demonstrates that a global increase in monomer levels upon stress fiber disassembly is sufficient to initiate NM2 filament assembly in the absence of actin dynamics.

**Fig. 4.**
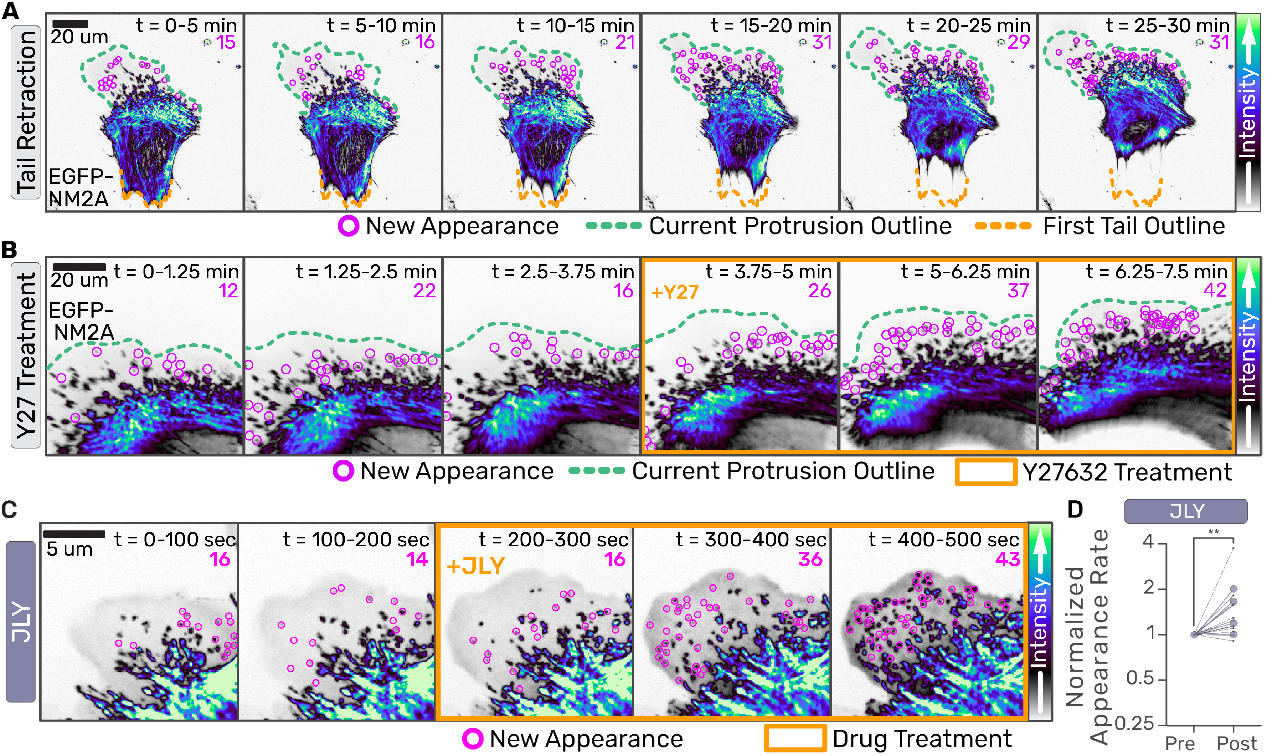
Globally elevating NM2 monomer availability initiates filament assembly. (a-c) Confocal z-stacks of primary EGFP-NM2A MEF cells were acquired. Images are sum intensity projections of Z-stacks and time-sum projections of 15 frames collected every 15 seconds in (a), 5 seconds in (b). Nascent NM2 filament appearances during that period are indicated with magenta circles and tallied in upper right corner. Green dotted line indicates current leading edge. Orange dotted line indicates original tail location (a) and orange box indicates frames treated with 10 uM Y-27632 (b). Scale bars = 20 um. (c) EGFP-NM2A sum projections were temporally-summed over 20 frames (100 seconds). Nascent NM2 filament appearances during that period are indicated with magenta circles and tallied in upper right corner. Three binned time periods pre- and -post (orange box) treatment are shown for JLY treatment. Scale bar = 5 um. (d) Nascent NM2 filament appearance events pre- and post-treatment were quantified, normalized to pre-treatment appearance events per minute, for individual cells (small black dots) and the mean of three independent experiments (large color circles). Intensity LUTs are indicated to the right of each panel. Wilcoxon matched-pairs signed rank test performed comparing each cell (n=12). p=0.0034

### Locally increasing NM2 monomer concentration initiates filament assembly

To directly test if artificially enhancing local NM2 monomer levels in a cell is sufficient to initiate NM2 filament assembly independent of upstream signaling, we engineered an improved light-inducible dimer (iLID) optogenetic system to optically recruit NM2 monomers to the cortex of migrating fibroblasts (28). We expressed a membrane-anchored LOV2-SsrA peptide in our Halo-tagged NM2A knock-in fibroblast cell line, along with a recruitable SspB-mApple-NM2A construct that can bind anchored SsrA upon blue light activation (Fig.5A). We then imaged the lamella while locally stimulating with blue light in a region devoid of NM2A filaments (Fig.5B, Movie 10). Within minutes, the photo-recruitable NM2A began accumulating in the stimulated region, followed shortly thereafter by the endogenous NM2A (Fig. 5C). Punctate filamentous structures containing a mixture of recruitable and endogenous NM2A continued to enrich and flow retrograde out of the stimulated region. These experiments reveal that locally increasing NM2 monomer concentration is sufficient to initiate filament formation, and that established NM2 filaments can enhance local filament assembly.

**Fig. 5.**
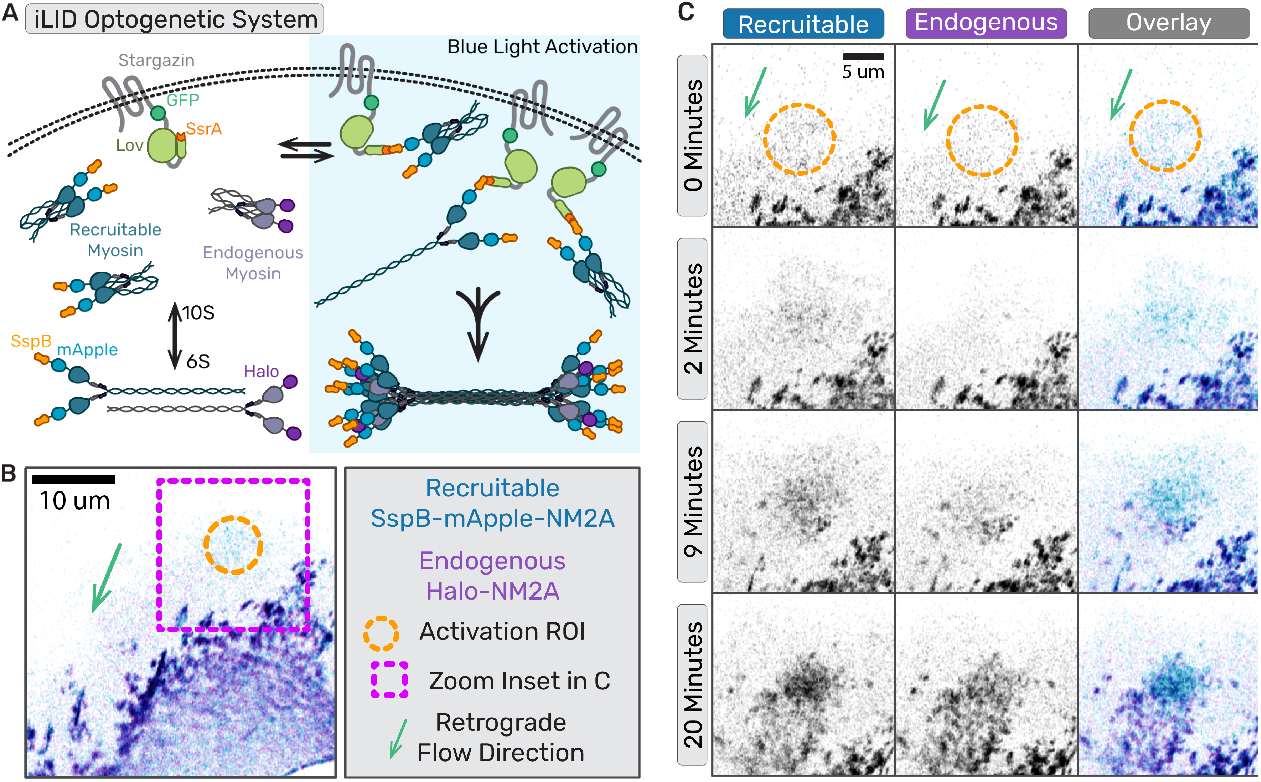
Locally concentrating NM2 monomer results in filament assembly. (a) Cartoon of iLID optogenetic system with photo-recruitable NM2 monomers expressed with endogenously tagged NM2. (b) Representative frame of optogenetic recruitment of NM2 monomers in cell. (b,c)Magenta box indicates the zoom inset region, orange circle marks the activated region, and green area shows the direction of retrograde flow. (c) Time series of optogenetic NM2 recruitment with recruitable NM2 in blue/grey and endogenous NM2 in purple/grey. Top panel is the recruitable NM2, middle panel is endogenous, and bottom panel is overlay of the two channels. Scale bar = 10 um.

### NM2 filaments established in the actin network are sites of enhanced filament assembly

Considering we could faithfully observe both NM2 filament initiating events and the enhancement of established NM2 filament clusters, we next sought to quantify their relative contributions to total lamellar filament assembly. Specifically, we asked if a new NM2 filament forms in the lamella, what is the likelihood that initiates a new cluster versus builds into an existing cluster. To do so, we adopted a molecular counting workflow using “standard candles” to build a standard curve of fluorescence and subsequently interpolate or extrapolate the number of NM2 monomers present in structures within the cell.

We used a membrane-anchored protein nanocage that self-assembles with 60 subunits when expressed in cells (Fig.6A) (29, 30). By using subunits with an EGFP on either one terminus (EGFP-60mer) or both termini (EGFP-120mer), we created two known standards. We created a third standard by expressing EGFP-Actin (Fig. 6B). Each standard candle was separately expressed in fibroblasts, where we segmented and quantified the fluorescent intensity of individual candles (Fig. 6C, Supp. Fig. 1A). We then created a standard curve by plotting the mean fluorescent intensities of each standard distribution as a function of the number of EGFP molecules present in the individual structures (Fig. 6D). Fitting a line to these data demonstrated a highly linear relationship between the fluorescence intensity of a structure and the known number of EGFP molecules present.

**Fig. 6.**
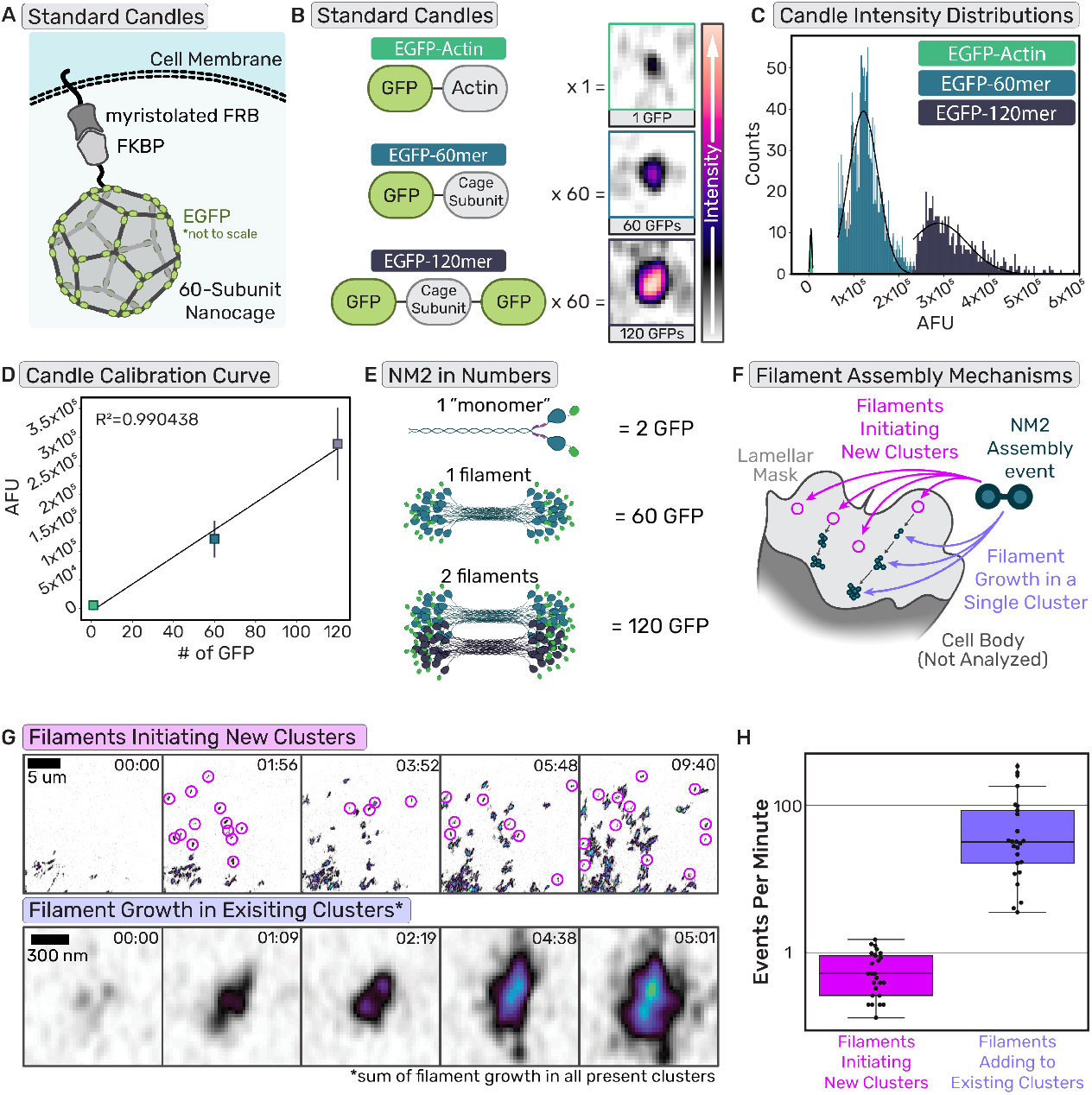
Established NM2 filaments are sites of enhanced filament assembly. (a) Cartoon of protein nanocage used as a known standard for the calibration curve. (b) Cartoons of the domains of the standard proteins used with the number of subunits per polymer and representative images of the standard in cells. Intensity scale bar on the right. (c) Intensity histograms of the known standards (d) Calibration curve generated from the intensity distributions of the standards and the number of subunits. n= 15 cells each candle, each experimental day. (e) Example cartoon of NM2 monomer, filament, and filaments tagged with EGFP with the number of EGFPs in those structures. (f) Cartoon delineating two different NM2 filament assembly mechanisms, initiation of new clusters or adding to existing clusters. (g)Example frames of cluster appearance tracking and NM2 filament growth tracking. Scale bar = 5 um; 300nm. (h) Quantification of cluster appearance rates compared to NM2 filament growth rates. n= 26 cells over 3 experiments

Using identical imaging settings, we then imaged endogenous EGFP-NM2A in fibroblasts from homozygotic knock-in mice. In these cells, every MHC 2A is tagged with an EGFP, and every NM2A monomer contains two EGFPs. *In vitro* studies and theoretical models demonstrate mature NM2A filaments consist of*∼* 30 monomers (5, 31, 32). Therefore, one mature NM2 filament would contain ~60 EGFP molecules while two mature filaments would contain *∼* 120 EGFPs (Fig. 6E), conveniently aligning with our known standards. Within lamellar regions (Fig. 6F), we quantified two parameters: (1) the number of nascent NM2 filament assembly events that initiate new clusters within a given time and (2) the fluorescent intensity increase for all NM2 adding to existing clusters in the same region during the same time (Fig. 6G-H, Movie 11). By converting the fluorescent intensity increase in all clusters to the number of NM2 filaments using our standard curve, we could directly compare the number of nascent filament assembly events (new clusters) to the number of filaments assembling into existing clusters. We find that assembling NM2 is *∼* 100 times more likely to incorporate into existing structures than to form nascent clusters in the lamella, demonstrating the dominant contribution of NM2-facilitated assembly to overall assembly.

### NM2 structures amplify to sub-resolution stacks before partitioning

Provided the importance of cluster growth to total lamellar NM2 assembly, and the derivation of clusters from a nascent NM2 filament, we sought to better define molecular mechanisms that enable addition of NM2 filaments to existing structures. Previous high resolution imaging studies observed the process by which a nascent NM2 filament grows in intensity before partitioning into multiple filamentous structures, a process that repeats sequentially to enhance cluster size as they mature into higher-order networks (Fig. 7A) (8, 9). Two non-mutually exclusive models were proposed in which a single mature NM2 filament is partitioned into two immature filaments (“Single Filament Partitioning”) or a mature filament recruits additional monomers/filaments to establish multiple filaments prior to partitioning (“Multi-Filament Partitioning”; Fig. 7B).

**Fig. 7.**
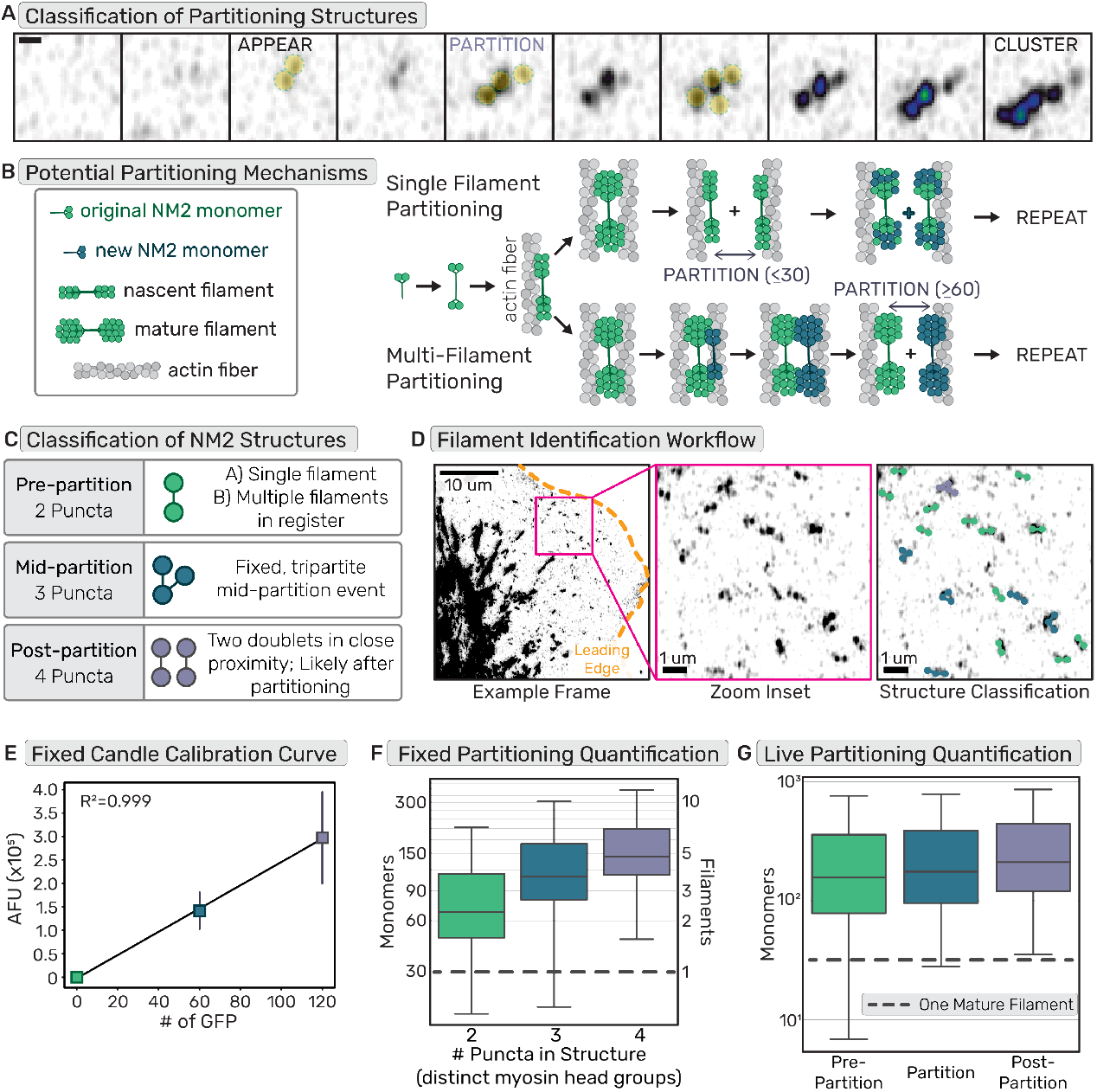
NM2 structures amplify to sub-resolution stacks before partitioning. (a) Example frames of EGFP-NM2A filament appearance, partitioning, and cluster growth. Scale bar = 300nm. (b) Cartoon depicting two non-exclusive mechanisms of NM2 partitioning, containing either single or multiple filaments. (c) Classification of single time-point or fixed frames of NM2 fluorescent structures grouping structures based on number of peaks/puncta and assigning them to stages of partitioning. (d) EGFP-NM2A sum projection example frame (left) with leading edge outlined in orange and zoom inset in magenta. Scale bar = 10um. Zoom inset (middle). Scale bar = 1um. Zoom inset overlay (right) displays detected NM2 filaments color coded into classes based on number of detected puncta in the fluorescent structures. Scale bar = 1um. (e) Example calibration curve for a fixed molecular counting data set. n= 15 cells each candle, each experimental day. (f) Fixed NM2 partitioning results with a grey dotted line to indicate a single mature NM2 filament. n= 45 cells over 3 experiments (g) Live NM2 partitioning results with a grey dotted line to indicate a single mature NM2 filament. n= 26 cells over 3 experiments

Using our molecular counting approach, we could now determine at which point during partitioning multiple NM2 filaments are present. We used super-resolution Airyscan imaging to resolve the number of GFP-tagged NM2 head groups within a structure. We classified two-puncta bipolar structures as pre-partitioning, three-puncta structures as mid-partitioning, and four puncta structures as post-partitioning (Fig. 7C-D, Movie 12). In the fixed imaging results, the identified two-puncta structures not only included structures just before partitioning, but also more nascent bipolar structures. Surprisingly, the vast majority of these two puncta structures already contained multiple NM2 filaments (Fig. 7E-F). This suggests a rapid amplification into sub-resolution filament stacks prior to spatial segregation of the NM2 filaments. The live results afforded the opportunity to isolate the exact frame prior to partitioning for each identified structure. Similar to the fixed results, most of the pre-partition or two puncta data contained multiple NM2 filaments. This number increased upon detectable partitioning and post-partitioning states (Fig. 7G). The higher number of NM2 monomers counted in the pre-partition data for live experiment compared to fixed results is likely due to our ability to identify partitioning events live and therefore filter out the more nascent two puncta structures that were not partitioning yet.

These data demonstrate that optically resolved two-puncta structures which have previously been identified as single NM2 filaments, are actually stacks of filaments in register. Partitioning is thus the separation of multiple NM2 filaments from one another, as opposed to the splitting of a single mature filament. Therefore, collectively, we suggest that cluster growth and overall NM2 assembly is highly driven by NM2-facilitated partitioning.

## Discussion

The demand for local force generation in numerous myosin 2-dependent processes dictates that NM2 filament assembly and amplification is dependent on its local environment. We propose here a mechanistic model where local myosin monomer availability enhances local NM2 filament assembly (Fig. 8). This myosin enrichment can be achieved strictly through dynamic reorganization of the actin cytoskeleton, including concentrating monomer in the lamella through collapse of leading edge protrusions (Fig. 8A-B) or globally elevating monomer availability via disassembly of actomyosin stress fibers in the rear of the cell during migration(Fig. 8A,C,J-K). At the heart of this model lies a reliance on cumulative actin-myosin and myosin-myosin interactions. While the kinetics of these many transient interactions will vary, the more local interactions there are, the more likely it is that sufficient monomers will dwell in a given area long enough to create a stable but immature NM2 filament. Once a nascent NM2 filament is established, it acts as a site of enhanced assembly, suggesting that myosin:myosin interactions are the main drivers of these amplification processes(Fig. 8D-E). The resulting clusters(Fig. 8F) of NM2 filaments continue to spatially segregate or partition(Fig. 8G), creating additional local assembly sites that further perpetuate the amplification and generation of force(Fig. 8H-I).

**Fig. 8.**
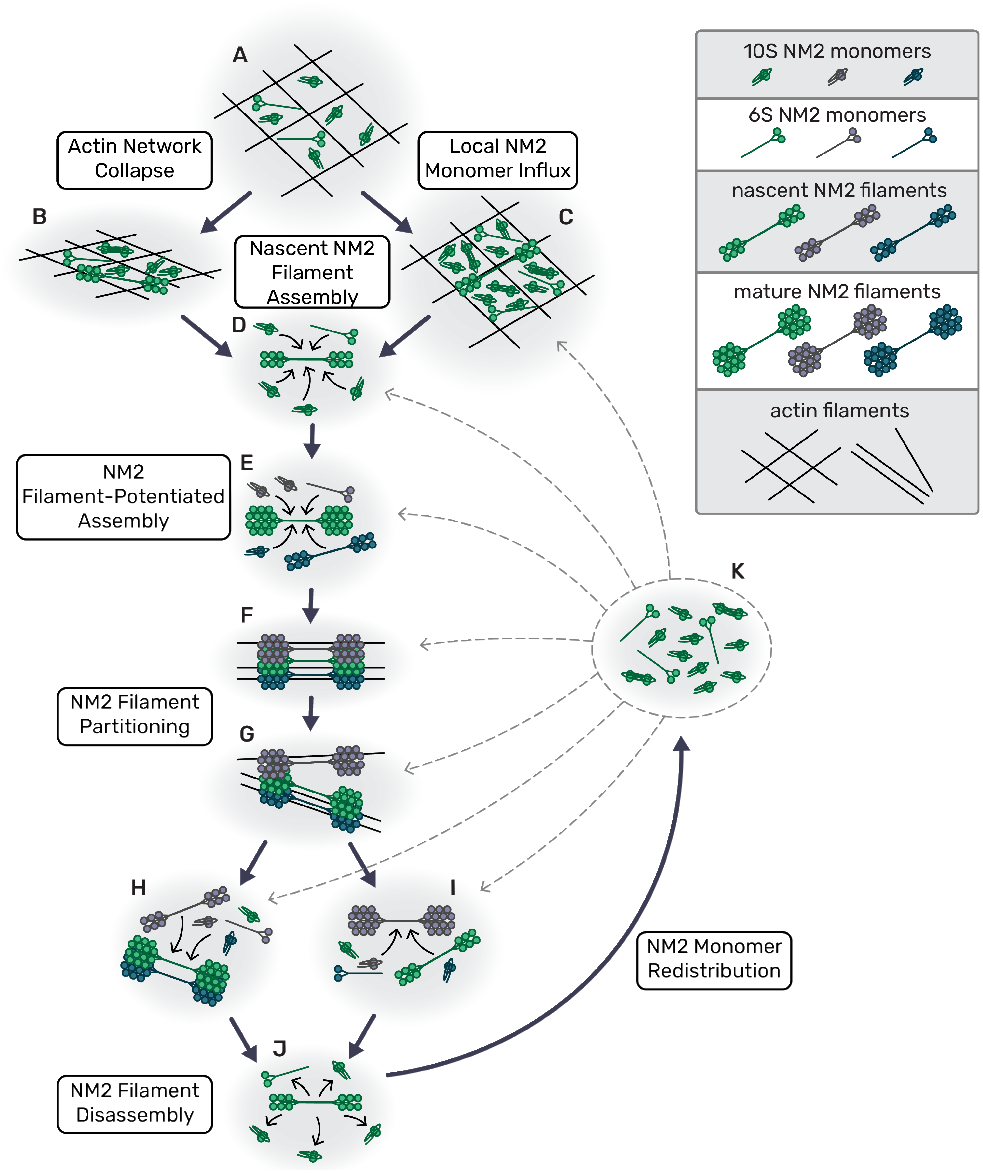
Model for NM2 filament assembly in cells. (a) Myosin monomers in actin network; large pore size in actin ‘net’. (b) Retraction collapses actin network, increases monomer density and potentiates NM2 filament assembly. (c) Myosin monomer concentration increases from NM2 filament turnover in other areas of the cell. (d) Local monomers add to nascent NM2 filament until it becomes a mature filament. (e) Established NM2 filament associates with local monomers and nascent filaments, potentiating assembly. (f) Sub-resolution NM2 filament stack/cluster continues to grow. (g) Sub-resolution NM2 filament stack/cluster partitions. (h,i) Separated, discrete myosin stacks and clusters continue to grow. (j) NM2 filament disassembles, releasing monomer. (k) Released NM2 monomers are redistributed throughout the cell, particularly aiding (c).

The first component of our model elicits a role for actin dynamics in dictating local myosin 2 assembly. Many previous studies have focused on the interaction between NM2 filaments and actin filaments, with myosin acting as either a motor or a crosslinker (33). In vitro and theoretical studies have shown that the presence of filamentous actin enhances NM2 filament assembly (7, 34). More recent cellular studies have reported actin-dependent roles in NM2 filament alignment, expansion, and partitioning (8, 9, 35). Here we reframe the perspective to focus on how actin dynamics impact myosin assembly. Our observations that leading edge and tail retractions precipitate NM2 filament formation suggest a role for the actin cytoskeleton in modulating myosin monomer concentration. Specifically, we speculate that leading edge retractions serve to locally concentrate myosin monomers by reducing the local actin pore size (36), similar to other models for actin-dependent restricted diffusion (37, 38). In addition to a reduced physical space, the increase in actin density would facilitate local NM2 retention by acting as a kinetic trap, providing a plethora of binding sites for free monomer to interact with. The ideal monomer to engage this actin is the 6S monomer transiently unfolded via phosphorylation of the RLC at Thr18/Ser19. However, even the inactive, folded monomer (10S) can bind actin, albeit with reduced affinity (39–41). Similarly, tail retraction and pharmacological disruption of the actomyosin cytoskeleton could spur additional NM2 filament assembly through release of NM2 monomer in filaments previously assembled in these structures. This sudden influx of NM2 monomers globally elevates the monomer concentration across the cell, facilitating nascent assembly by increasing the likelihood of NM2 monomer interactions. In each case the dynamics of the actin network serves to modulate NM2 concentrations that drive assembly of NM2 filaments.

The second overarching component of our model elicits myosin:myosin interactions. In this regard, monomer-monomer, monomer-filament, and filament-filament interactions should be considered. It is known that multiple tail interactions between several monomers within a NM2 filament help to stabilize the structure (31). While the tail interactions surely dominate, numerous other interactions have been identified within the myosin holoenzyme (head-head, head-tail, light chain-tail, etc.) (42). It is quite possible that these low affinity interactions also occur between NM2 monomers and filaments to contribute to the local enrichment of myosin. Additionally, NM2 filament stacks and clusters, concatenation and partitioning have all been reported through a combination of in vitro and cellular studies (8, 9, 43). This indicates myosin-myosin interactions beyond those occurring between monomers within a filament. Based on FRAP studies, the half-time of recovery is typically within 10s of seconds, indicating rapid NM2 filament exchange kinetics (44, 45). We hypothesize that once a nascent NM2 filament is established, transient interactions and rapid exchange kinetics enrich the local monomer concentration to increase the probability of additional filament assembly events (46). We speculate this effective diffusion trap is the basis for the myosin-facilitated assembly that we observe in our data. Importantly, this model is not mutually exclusive with the potential for a nascent NM2 filament to enhance filament assembly via mechanosensitive feedback systems that alter local actin to favor assembly or initiate signaling events that lead to canonical myosin activation via RLC phosphorylation (16, 47–49). Indeed, our observation of active Rho flanking established myosin clusters is supportive of parallel mechanisms to amplify myosin filaments.

We propose that these mechanisms contribute to the rapid assembly and amplification of NM2 filaments to efficiently produce physiological levels of contraction in polarized migration. While it is most straightforward to experimentally observe these myosin dynamics in the lamellar regions of migrating cells, we believe this to be a universal mechanism of rapidly building contractility. Two additional areas of biology that have clear evidence for, and use of, rapid myosin filament assembly and amplification are the contractile ring (50) and adherens junction maturation (51). In both contexts, higher-order myosin networks are observed and must develop rapidly to achieve the requisite contractility. The mechanisms that we outline here could help to drive the NM2 filament amplification in these dense areas. While additional work will be needed to confirm these mechanisms at work in dense regions, it is clear that in addition to biochemical regulation, myosin filament assembly and amplification are sensitive to biophysical constraints.

## Materials and Methods

### Cell culture and transfection

Primary MEFs previously isolated (9) from EGFP NM2A knock-in mice [PubMed: 21908426] were used within 10 passages. The JR20 cell line of immortalized MEFs (13) were cultured within 15 passages. HEK293T cells were cultured to prepare lentivirus. Cells were cultured in Dulbecco’s modified Eagle’s Medium (Corning #10-013-CV) supplemented with 10% FBEssence (Avantor Seradigm #10803-034) and 1X antibiotic-antimycotic (Corning #30-004-CI) at 37°C in 5% CO2. All cells were grown on uncoated plastic tissue culture dishes and plated on #1.5 coverslips (Cellvis #C8-1.5H-N) pre-coated overnight at 4°C with 10µg/mL fibronectin [Fisher Scientific #FC01010MG] then rinsed with PBS before plating. Live-imaging experiments were conducted with Leibovit’s L-15 medium without phenol red (Gibco #21083027) supplemented with 10% FBEssence.

Cells were transfected for CRISPR knock-in, lentivirus preparation, and the calcium and rho biosensor experiments using the LipoD293 (SignaGen #SL100668) system.[ADD DNA CONC, LIPOD AMOUNT, CELL AMOUNT] Cells were transfected for molecular counting experiments using the Neon Electroporation system [ThermoFischer Scientific] with 2 × 20msec pulses of 1350V and 5*µ*g plasmid DNA into 400k cells in a 100uL reaction. For calcium activity experiments, cells were transfected with GCaMP7s (Addgene Plasmid #104463)(14), plated 24-48 hours post-transfection and 12-24 hours before imaging, and then cells of average brightness were imaged. For RhoA activity experiments, cells were transfected with GFP-RhoA-AHPH (AKA GFP-RhoBio) (Addgene Plasmid #68026)(15) with the same approach as described for the calcium experiments. Cells transfected with the protein nanocages for the molecular counting experiments were transfected with mem-EGFP-60mer or mem-EGFP-120mer, plated 48 hours post-transfection and 18-24 hours before imaging. Cells transfected with EGFP-Actin (Addgene plasmid #31502) (52) for the molecular counting experiments were transfected 4-6 hours prior to imaging and plated immediately to achieve extremely low expression for single molecule identification. Primary MEFs used for the long term cell migration and JLY experiments were infected with viral media from HEK cells transfected with 3x-mScarlet-FTractin (Addgene Plasmid #112960)(53). Once stably expressing the 3x-mScarlet-FTractin, the cell line was re-stored and used within 10 passages. For the optogenetics experiment, a stable cell line was created using pLV-Stargazin-mTurquoise2-iLID (Addgene Plasmid #161001) (54) lentivirus and then transiently transfected with SspB-mApple-NM2A using LipoD293 transfection system. Cells were plated 24-48 hours post transfection and 12-24 hours before imaging.

Inhibitors were used at the following concentrations: Y27631 (EMD Millipore #68801), 10*µ*M; LatrunculinB (EMD Millipore #428020), 1.25*µ*µM; Jasplakinolide (EMD Millipore #420127), 2*µ*M. Drug treatments were prepared at a 2X solution in L-15 imaging media and then added to the wells at a 1:1 dilution while imaging. All cell lines were tested for mycoplasma once a month.

### Generation of CRISPR knock in cell lines

Halo-NM2A and mScarlet-NM2A knock-in cells were derived from JR20 parental fibroblasts using CRISPR/Cas9. We generated pSpCas9(BB)-2A-Puro (PX459) V2.0 (Addgene Plasmid #62988) with target sequence AAACTTCATCAATAACCCGC using established protocols To generate donor plasmids, pUC57 was digested with EcoR1 and Stu1 and purified. A four piece Gibson assembly was then performed with three gBlocks (IDT): 1) a 794 bp 5’ HDR of genomic sequence immediate upstream of the endogenous start codon, 2) mScarlet fluorophore or HaloTag with an 18 amino acid GS-rich linker, 3) a 802 bp 3’ HDR of genomic sequence immediately downstream of the endogenous start codon with silent PAM site mutation. JR20 were transfected with donor and target-Cas9 plasmids, single-cell sorted at 5-10 days post-transfection. Individual clones were evaluated for knock-in via western blotting and microscopy. Clones used in this study include Halo-NM2A clone 2 (H2A2) and mScarlet-NM2A clone 3 (S2A3).

### Molecular cloning

To engineer photo-recruitable NM2A, we introduced an SspB upstream of mApple in pmApple-NM2A. After digesting pmApple-NM2A with Age1, an SspB PCR product with flanking HDR arms was introduced via Gibson cloning in frame with mApple-NM2A.

### Lentiviral Stable Cell Lines

HEK293T were transfected using LipoD293 (Signagen #SL100668) and the accompanying lentivirus generation transfection protocol. Briefly, cells were plated in a 6cm dish and grown to 80-90% confluence. Approximately one hour before transfection, media was changed on the cells. Transfection complex was created with LipoD293, packaging plasmic psPax (Addgene Plasmid #12260) and envelope plasmid PmD2.G (Addgene Plasmid #12259), and lentiviral construct and added dropwise to the dish. Media was changed at 24 hours post transfection and collected at 48 and 72 hours. Viral media was spun down at 1000g for 5 minutes and then filtered with a 0.45*µ*m filter. A 50% confluent 6cm of cells were then infected with 48 hour viral media. Viral media was removed after 24 hours and cells stored for future use after 3-5 days.

psPAX2 was a gift from Didier Trono (Addgene plasmid # 12260; http://n2t.net/addgene:12260; RRID:Addgene_12260). pMD2.G was a gift from Didier Trono (Addgene plasmid # 12259; http://n2t.net/addgene:12259; RRID:Addgene_12259)

### Imaging

*Calcium and Rho:* Calcium and Rho imaging was performed on a Zeiss 880 Airyscan with a 63x 1.4 NA objective in the “Airyscan Fast” acquisition mode. Time lapse images of NM2 and calcium or rho biosensors were conducted at a 2.5 second or 10 second frame interval, respectively. Long term migration imaging was performed on a 3i confocal spinning disk with a 40x 1.3 NA objective. Time lapse images of NM2 (488) and actin (561), at 30% laser and 50msec exposure per channel. Between 12 and 15 positions were acquired every minute for eight hours. Zeiss definite focus.2 was used to focus between timepoints and positions.

*JL and JLY:* JLY experiments were performed on a 3i confocal spinning disk with a 63x 1.4 NA objective. Time lapse images of NM2 (488) and actin (561), at 30% laser and 50msec exposure per channel. Four positions were acquired every five seconds for ten minutes. Zeiss definite focus.2 was used to focus between timepoints and positions. After 5 minutes, drug cocktails prepared at a 2X concentration were added 1:1 with the media in the well while imaging to ensure rapid treatment. Cells were imaged until they ripped apart from the drug treatment and then an equal number of frames before and after treatment were used for analysis.

*Molecular counting:* For all molecular counting experiments, standard candle controls and EGFP-NM2A imaging were performed each day within 6 hrs of each other using identical laser and acquisition settings. Live molecular counting experiments were performed on a Zeiss 880 Airyscan with a 63x 1.4 NA objective in the “Airyscan Fast” acquisition mode. Argon laser at 0.9% power with 0.83*µ*sec pixel dwell, a 4x line averaging was used for all images acquired for the standard curve and live imaging of NM2 to be counted. Time lapse images of NM2 were conducted with identical laser power, pixel dwell, and averaging at a 5 second frame interval.

Fixed molecular counting experiments were performed on a Zeiss 880 Airyscan with a 100x 1.4 NA objective in the “Airyscan Fast” acquisition mode. Argon laser at 9% power with 0.98*µ*sec pixel dwell, a 4x line averaging was used for all images acquired for the standard curve and live imaging of NM2 to be counted. Time lapse images of NM2 were conducted with identical laser power, pixel dwell, and averaging at a 5 second frame interval.

### Correlative Light And Electron Microscopy

Round 25mm coverslips were squeaky-cleaned (56) and plasma cleaned before coating with 20*µ*g/mL Human Plasma Fibronectin (EMD Millipore #FC010) at 37C for 1 hour. After fibronectin coating and subsequent PBS washes, PDMS strips approximately 5mm in width and 25mm on length, were placed to bisect the circular coverslip. Cells were plated at 75% confluence and incubated overnight. 20-24 hours after plating, PDMS strips were removed and the media changed twice to remove any lifted cells. Cell unroofing was performed 12-18 hours after PDMS strip removal as described previously (22, 23). Briefly, coverslips individually taped onto a 6cm petri dish and covered with PBS. Then, they were rinsed with intracellular buffer (70mM KCL, 30mM HEPES maintained at pH 7.4 with KOH, 5mM MgCL2, 3mM EGTA) and then cell edges ‘glued’ down with a 30 second treatment of 0.08% Poly-L-Lysine in intracellular buffer. Then cells were manually unroofed by spraying 1 mL of 3% paraformaldehyde (Electron Microscopy Sciences #15710) and 1% glutaraldehyde (Electron Microscopy Sciences #16216) in intracellular buffer through a 25 gauge needle along the line of the PDMS ‘wound’ from a distance of 1cm. Fresh fixative was added to fully submerge the coverslips to fix for 30 minutes at room temperature. Coverslips were then rinsed with PBS and then removed from the petri dishes to incubate in 1:50 phalloidin-555 (Thermo Fisher #A34055) face down on parafilm for 30 minutes at RT. Immediately after phalloidin staining, coverslips were carefully flipped over, avoiding any sliding, and placed in a PBS wash. The coverslips were then mounted in a magnetic chamber in PBS and immediately imaged on the Airyscan 880. Cells that appeared unroofed based on phalloidin intensity were selected within a 1mm diameter and the coverslip was marked for that region with a diamond tip objective. Within the marked region, NM2 and actin were imaged at 63x 1.4 NA oil in “SR” mode and then 40x 1.3 NA oil tilescan in “Fast” mode. Coverslips were then imaged using EVOS phase contrast microscope at 20x, 10x, and 4x. Images were then organized to create a map back to the same imaged ROI after platinum replica preparation. After fluorescence imaging, coverslips were placed in fixative, flipped over onto a glass slide. The coverlsips were immobilized with epoxy resin, sealed with VALAP, and shipped overnight at 4°C to the Taraska lab at NIH for the PREM workflow.

Platinum replica sample preparation was performed as described in (22, 23). Briefly, coverslips were placed in 0.1% tannic acid for 20 minutes, rinsed 4x in water, 0.1% uranyl acetate for 20 minutes, rinsed 2x in water, and then dehydrated gradually with increasing concentrations of ethanol (15, 30, 50, 70, 80, 90% up to being rinsed 3x in 100% ethanol) prior to critical point drying (Tousimis 895). After critical point drying, the coverslips were trimmed down with a diamond scriber. Samples were rotary coated with a 2-3 nm coat of platinum-carbon at a 17 degree angle, then 5-6 nm carbon at a 90 degree angle (RMC 9010). The coverslips were imaged with 20x phase contrast light microscopy to find the cells that were previously mapped during fluorescence imaging. The coverslip was then placed face up on the air/water interface of 5% hydrofluoric acid until the coverslip dropped into solution leaving the platinum replica sitting on surface. The replica was rinsed with water and lifted with a 4 mm circular loop onto a formvar/carbon coated 75-mesh copper grid (Ted Pella 01802-F). The grid was again imaged with 20x phase contrast light microscopy to confirm that replica transfer went smoothly and identify the area of interest on the grid. Transmission electron microscopy (TEM) was performed with montaging on a FEI Tecnai T12 equipped a Gatan Rio-9 camera and SerialEM freeware (57). TEM microscope was maintained by Haotian Lei and Yanxiang Cui as part of the NIDDK cryo-EM facility at the NIH Bethesda campus.

Correlation of fluorescence and platinum replica electron micrographs was performed using Matlab correlation software (23). Major structures in the actin channel (e.g. large bundles or unique branching) were used to correlate images, with a minimum of 20 points used to correlate.

### Image Analysis

*Calcium, Rho, Y27, and Tail Release:* Qualitative analysis of NM2 filament appearance was conducted in FIJI by making sum time projections for every 10 frames. Kymographs were generated in FIJI using the KymoResliceWide plugin with a line width of 11.

*JL and JLY:* Qualitative analysis of NM2 filament appearance was conducted in FIJI by making sum time projections for every 10 frames. Quantification of NM2 filament appearance for JLY analysis was performed manually in FIJI by identifying puncta appearance in the lamella over the course of all of the frames. Average appearance rates for all frames acquired after drug treatment and the same number of frames pre-treatment and normalized to the pre-treatment rate for each cell to compare paired, before and after treatment conditions.

*Molecular counting:* Image analysis was performed using custom written python analysis software (https://github.com/m-a-q or https://github.com/OakesLab). Images were sum-projected in Z and then cell masks generated for each image and each frame for time series. For candle images, single frames were analyzed and local peaks (or puncta) identified within the cell mask. The intensity of a 14 by 14 pixel box around each peak was used to filter out any candles that were side by side. The intensity of the top of bottom z slices where the peaks are identified in X and Y was used to filter out candles that were not fully within the Z range that was acquired. Once unsuitable peaks were filtered out, the remaining list of peak positions were used to quantify the sum intensity of a 14 by 14 pixel box around the peak coordinates. Candle intensities were then plotted in histograms and then plotted by the number of GFPs present in the molecules. The intensity of a single GFP was found from linear regression analysis.

Images of NM2 were analyzed using trackpy (https://github.com/soft-matter/trackpy) (58) to first identify NM2 filament clusters as they appear and flow back with retrograde flow. A 35 by 35 pixel box around the particle centroid was summed to quantify the fluorescent intensity, and then divided by the intensity of a single GFP, then divided by 2, to get the number of NM2 monomers present in the structure at any given frame. This was compared to the rate of track appearance from the same trackpy analysis. For partitioning analysis, Individual tracks were then further tracked to identify local peaks within the growing cluster to count the number of NM2 head groups that could be resolved. Based on the number of resolvable head groups, we determined the partitioning state of the structure.

### Statistical analysis

To compare the number of NM2 filament appearance rates (Fig. 2C, Fig. 4D), we used a Wilcoxon matched-pairs signed rank test performed comparing ‘pre’ and ‘post’ frames for each cell. Statistical analysis was performed using Prism (GraphPad). p-values less than 0.05 were considered significant.

Intensity histograms (Fig 6.c) were fits to gaussians and linear regression analysis used on the calibration curves (Fig. 6D, Fig. 7E) to determine the value of a single GFP. Statistical analysis was performed using the stats module from scipy.

## Data availability

The data that support the findings of this study are available upon reasonable request from the corresponding author [Jordan Beach].

## Acknowledgements

We thank the rest of the Beach and Oakes laboratories at the Loyola University Chicago, Maywood, IL for many helpful discussions. Research reported in this publication was supported by the Maximizing Investigators’ Research Award (MIRA R35) from the National Institute of General Medical Sciences (NIGMS) of the National Institutes of Health (NIH) #R35GM138183 to JRB, National Science Foundation CAREER Award #2000554 and NIH National Institute of Allergy and Infectious Disease (NIAID) (Award # P01-AI02851) to PWO, and the National Science Foundation Graduate Research Fellowship (NSF GRFP) #DGE-1842190 to MAQ.

## Author contributions

MAQ, PWO and JRB designed the research. MAQ, KAS, HW, HP, and JRB performed experiments. MAQ, PWO, and JRB analyzed data. JDR and JEB contributed critical cell lines. FK, JWT, and MA contributed to experimental techniques. MAQ, PWO and JRB wrote the manuscript with critical feedback from co-authors.

## Supplementary Information

**Movie 1 - NM2 Filament appearance tracking**. Sum intensity Z-projection of time lapse of EGFP-NM2A knockin cell. Lamellar protrusion imaged with a 5 second frame rate. The leading edge is in the top right corner with retrograde flow to the bottom left. Magenta circles mark NM2 filaments in the first frame of appearance. Scale bar = 5 um. Time = mm:ss.

**Movie 2 - Leading edge retraction precedes NM2 filament appearance**. Sum intensity Z-projection of time lapse of EGFP-NM2A knockin cell. Lamellar protrusion imaged with a 1 second frame rate. The leading edge is in the top right corner with retrograde flow to the bottom left. Scale bar = 5 um. Time = mm:ss.

**Movie 3 - JL treatment stalls actin dynamics**. Sum intensity projection of Z-stack time lapse of 3x-mScarlet-FTractin tagged lamellar protrusion with a 5 second frame rate. The leading edge is in the top right corner with retrograde flow to the bottom left. Scale bar = 5 um. Time = mm:ss.

**Movie 4 - DMSO treatment with NM2 appearance**. Sum intensity Z-projection of time lapse of EGFP-NM2A knockin cell expressing 3x-mScarlet-FTractin. Lamellar protrusion imaged with a 5 second frame rate. The leading edge is in the top right corner with retrograde flow to the bottom left. Actin is in grey LUT and NM2A in blue LUT. Drug treatment indicated in orange text at the first frame of treatment.Scale bar = 5 um. Time = mm:ss.

**Movie 5 - JL treatment with NM2 appearance**. Sum intensity Z-projection of time lapse of EGFP-NM2A knockin cell expressing 3x-mScarlet-FTractin. Lamellar protrusion imaged with a 5 second frame rate. The leading edge is in the top right corner with retrograde flow to the bottom left. Actin is in grey LUT and NM2A in blue LUT. Drug treatment indicated in orange text at the first frame of treatment. Scale bar = 5 um. Time = mm:ss.

**Movie 6 - Actomyosin dynamics during long term cell migration**. EGFP-NM2A knockin cells expressing 3x-mScarlet- FTractin imaged every minute for 8 hours. Actin is in grey LUT and NM2A in blue LUT. Scale bar = 50 um. Time = hh:mm.

**Movie 7 - Tail retraction precedes NM2 filament appearances**. Sum intensity Z-projection of time lapse of EGFP-NM2A knockin cells imaged with a 20 second frame rate. Initial tail outline in orange and initial protrusion outline in green. Scale bar = 20 um. Time = mm:ss.

**Movie 8 - Y27 treatment results in increased NM2 filament appearances**. Time lapse of EGFP-NM2A knockin cell with a 5 second frame rate. Drug treatment indicated in orange text at the first frame of treatment. Scale bar = 50 um. Time = hh:mm.

**Movie 9 - JLY Treatment with NM2 Appearance**. Sum intensity Z-projection of time lapse of EGFP-NM2A knockin cell expressing 3x-mScarlet-FTractin. Lamellar protrusion imaged with a 5 second frame rate. The leading edge is in the top right corner with retrograde flow to the bottom left. Actin is in grey LUT and NM2A in blue LUT. Drug treatment indicated in orange text at the first frame of treatment. Scale bar = 5 um. Time = mm:ss.

**Movie 10 - NM2 Monomer Recruitment Jumpstarts NM2 Filament Assembly**. Sum intensity Z-projection time lapse of Halo-NM2A knockin cell expressing GFP-Stargazin-Lov-SsrA and SspB-mApple-NM2A. Lamellar protrusion imaged with a 10 second frame rate. The leading edge is in the top right corner with retrograde flow to the bottom left. Recruitable (SspB-mApple) NM2 shown in the left panel, endogenous (Halo-646-NM2) in the middle panel, and an overlay with recruitable in blue and endogenous in purple in the right panel. Orange circle marks the activation ROI that was stimulated after each acquisition frame. Scale bar = 5 um. Time = mm:ss.

**Movie 11 - NM2 Filament mediated amplification**. Sum intensity Z-projection of time lapse of EGFP-NM2A knockin cell. Lamellar protrusion imaged with a 5 second frame rate. The leading edge is in the top right corner with retrograde flow to the bottom left. Scale bar = 5 um. Time = mm:ss.

**Movie 12 - NM2 Filament Partitioning**. TIRF-SIM time lapse of EGFP-NM2A knockin cell with a 2 second frame rate. Green circles indicate the first frame with 2-puncta *∼* 300 nm apart, or pre-partitioning. Blue circles indicate the first frame with 3-puncta 3*∼* 00 nm apart, or mid-partitioning. Purple circles indicate the first frame with 4-puncta *∼* 300 nm apart, or post-partitioning. Scale bar = 300 nm. Time = mm:ss.

---

**Fig. S1.**
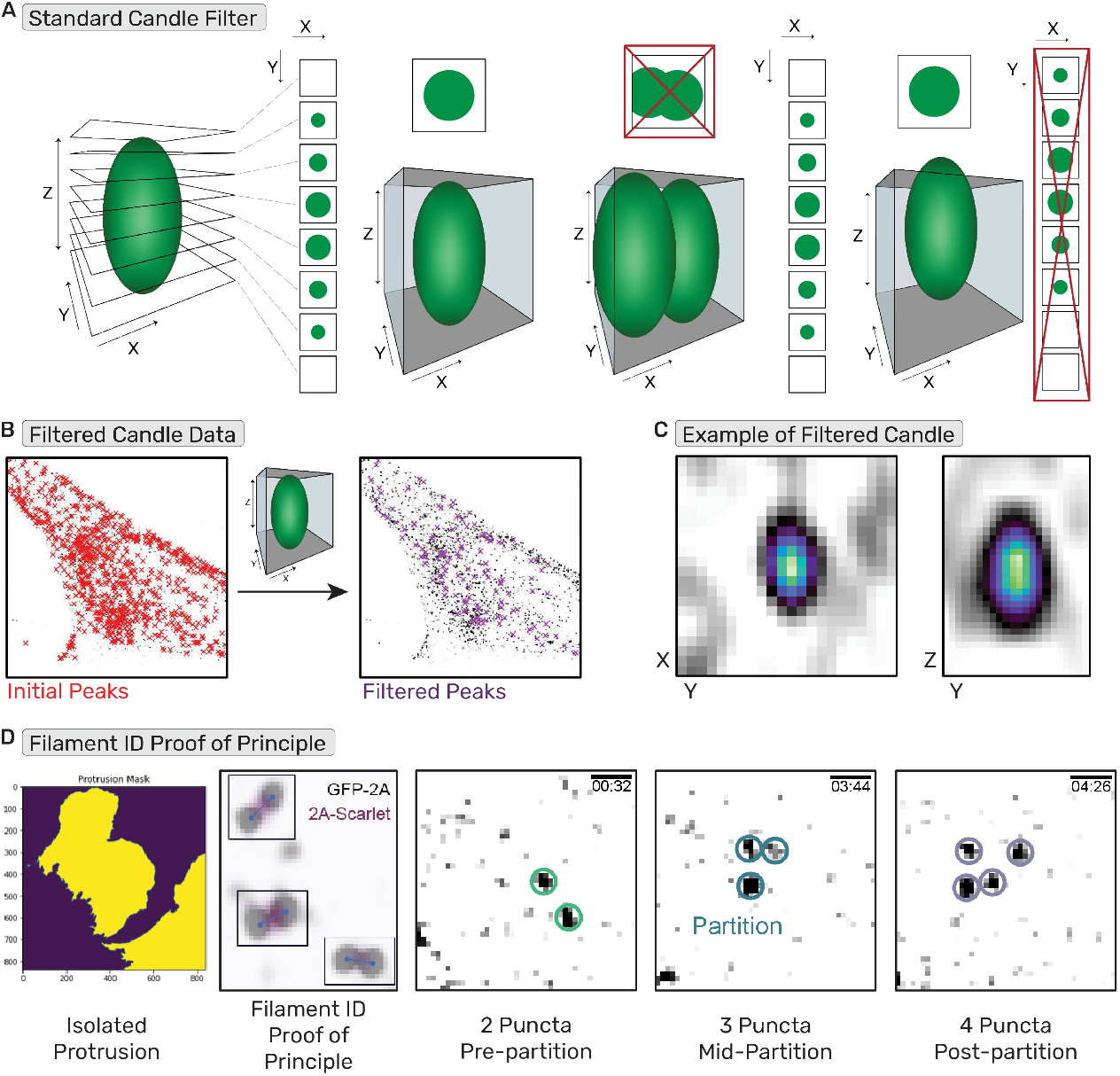
Molecular Counting Workflow. (a) Depiction of filter used to identify candles unsuitable for analysis (i.e. too close to another candle or partially out of Z-range). (b) Example of standard candle image before and after filter. (c) Example image of passable filter in XY and ZY. (d) Depiction of NM2 filament identification workflow with (1) lamella isolation, (2) filament identification by puncta ^*∼*^300nm apart with proof of principle NM2-tail tag to confirm filament, and (3) examples of the 2-puncta, 3-puncta, and 4-puncta identifications.

